## Supplementary figures and images for "*Salmonella* exploits host- and bacterial-derived β-alanine for replication inside host macrophages"

### Figure 1_figure supplement 1

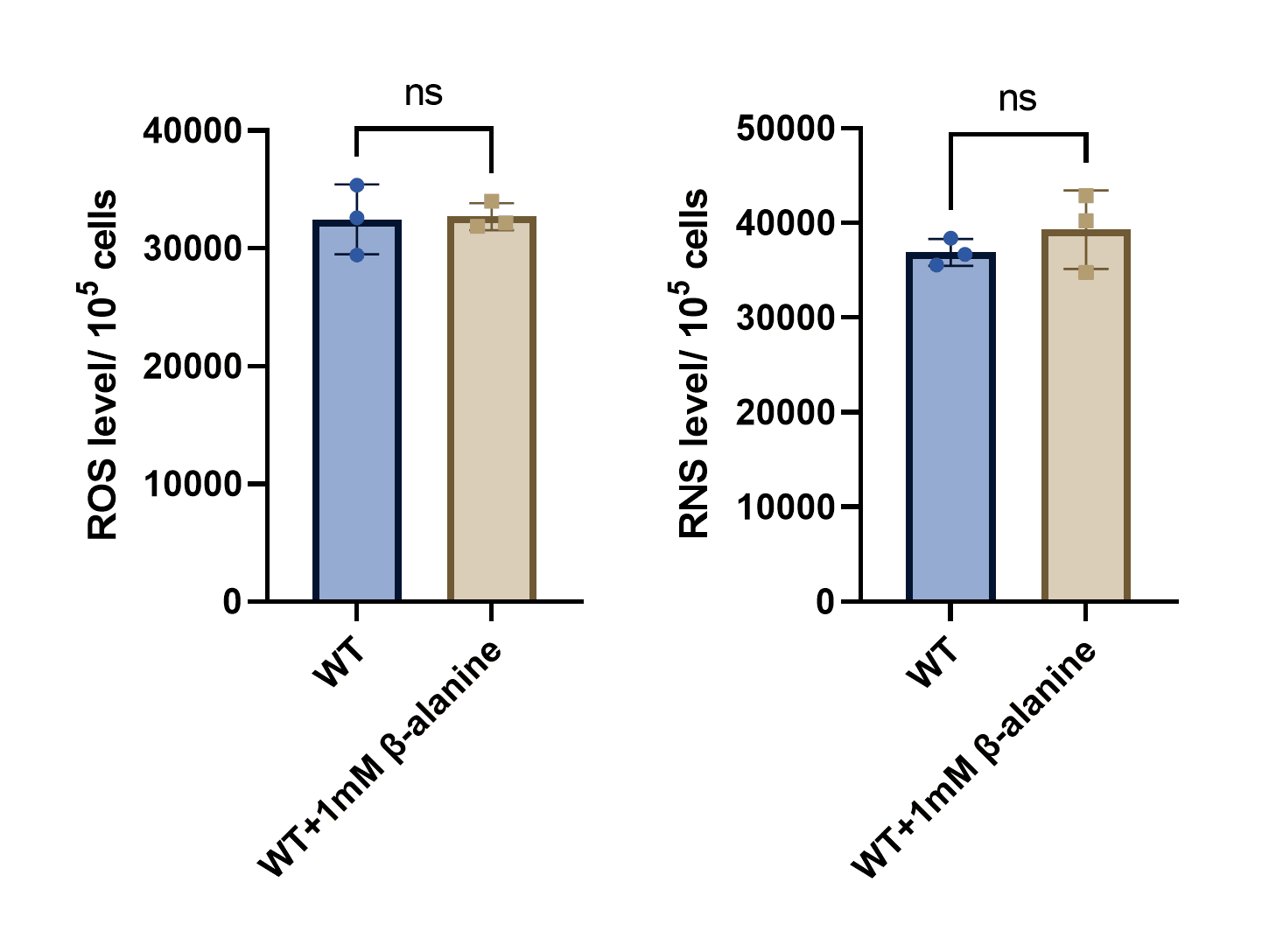

### Figure 1_figure supplement 2

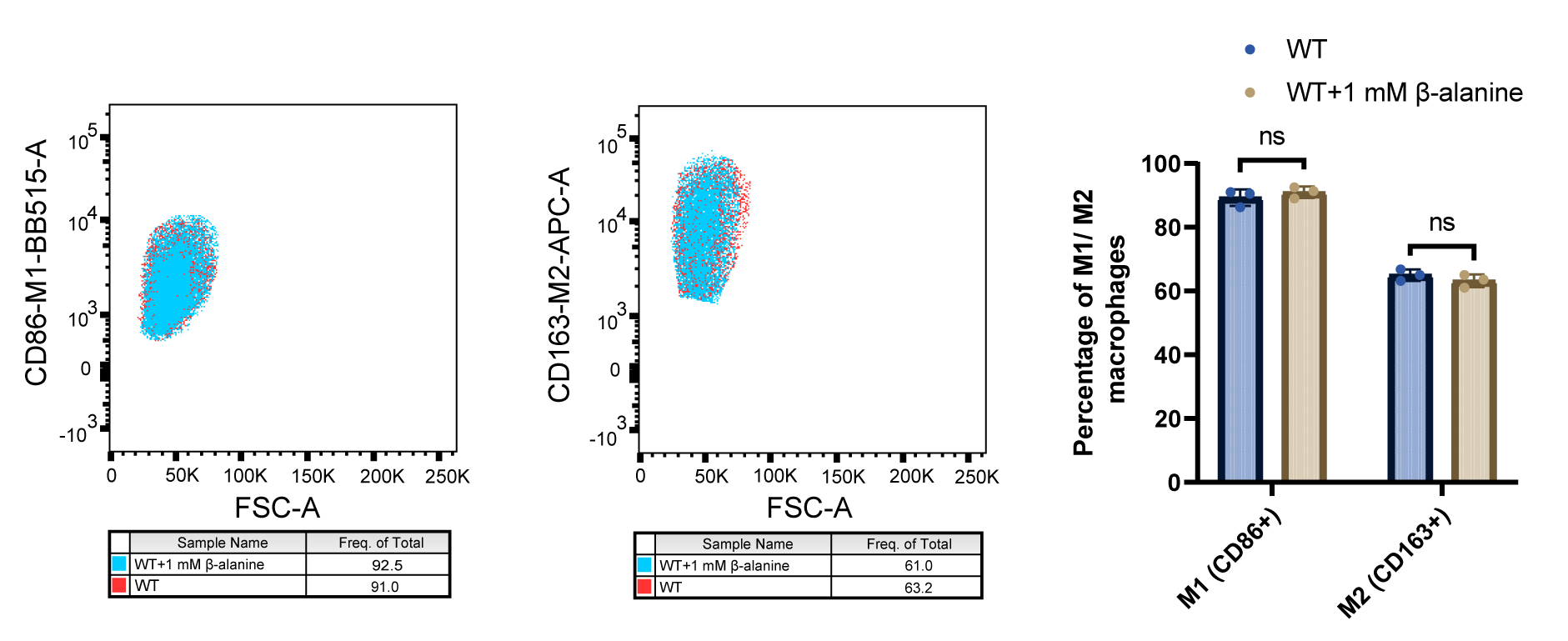

### Figure 1_figure supplement 3

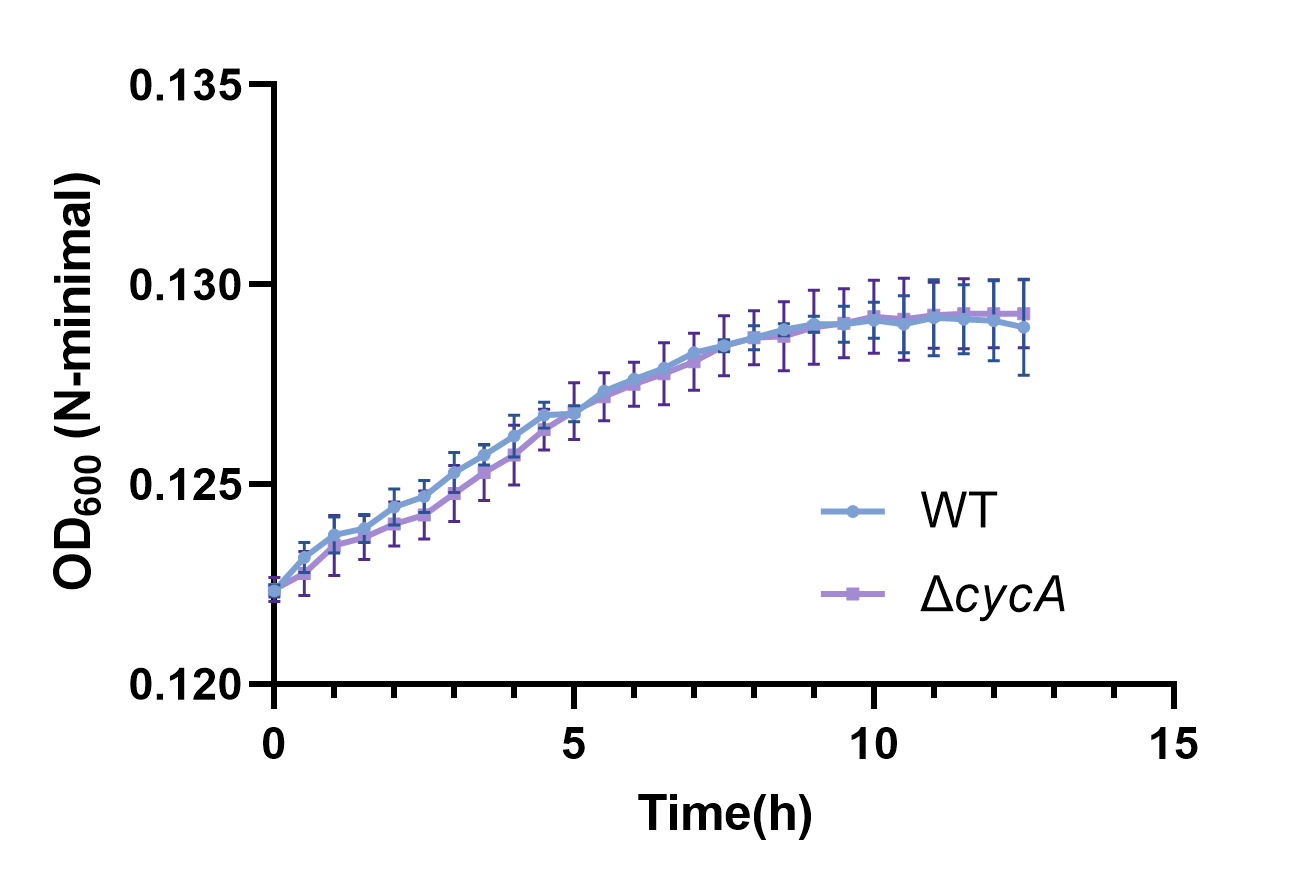

### Figure 1_figure supplement 4

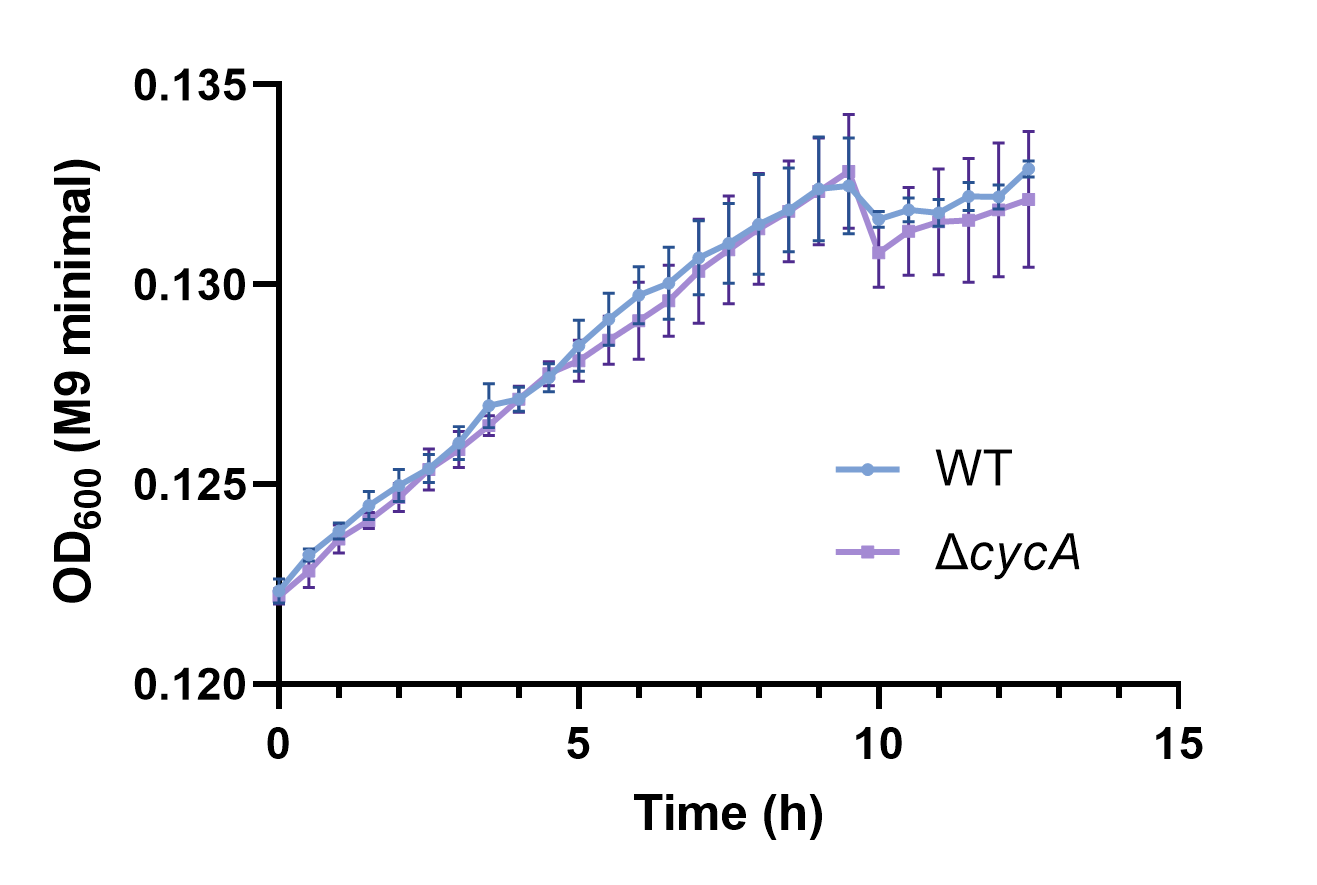

### Figure 1_figure supplement 5

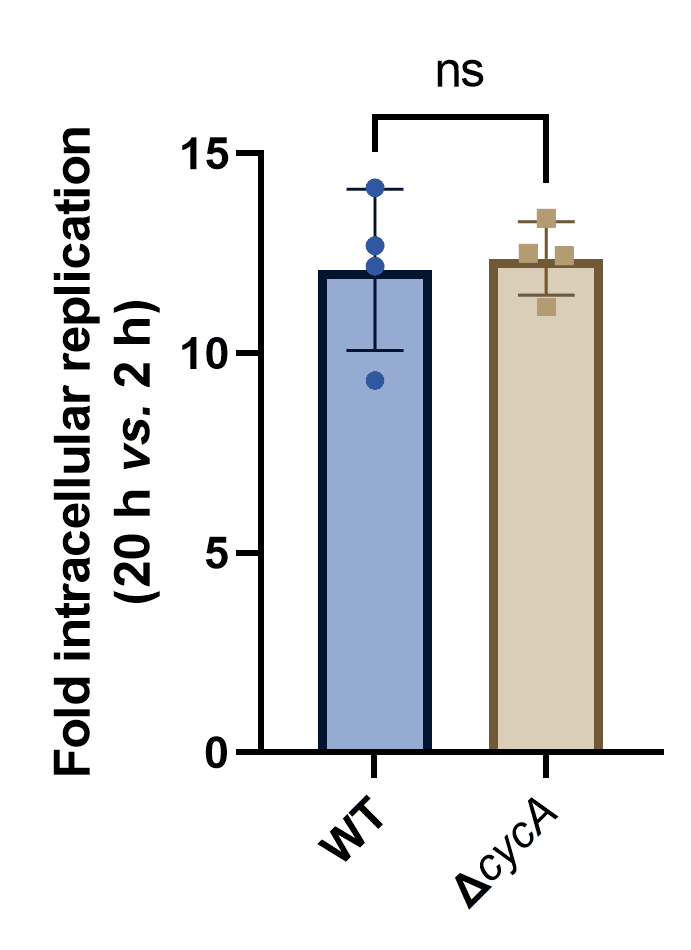

### Figure 1_figure supplement 6

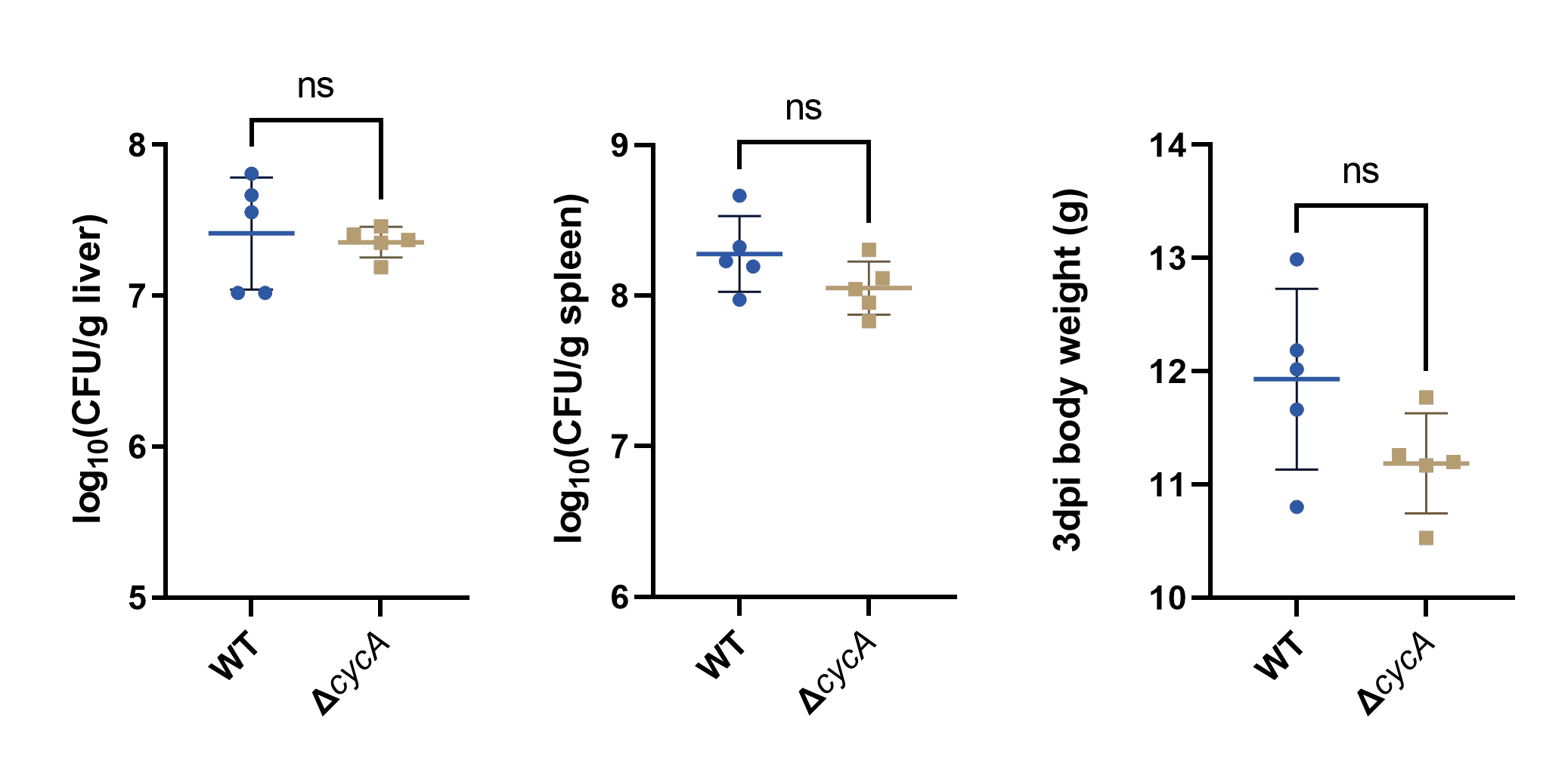

### Figure 1_figure supplement 7

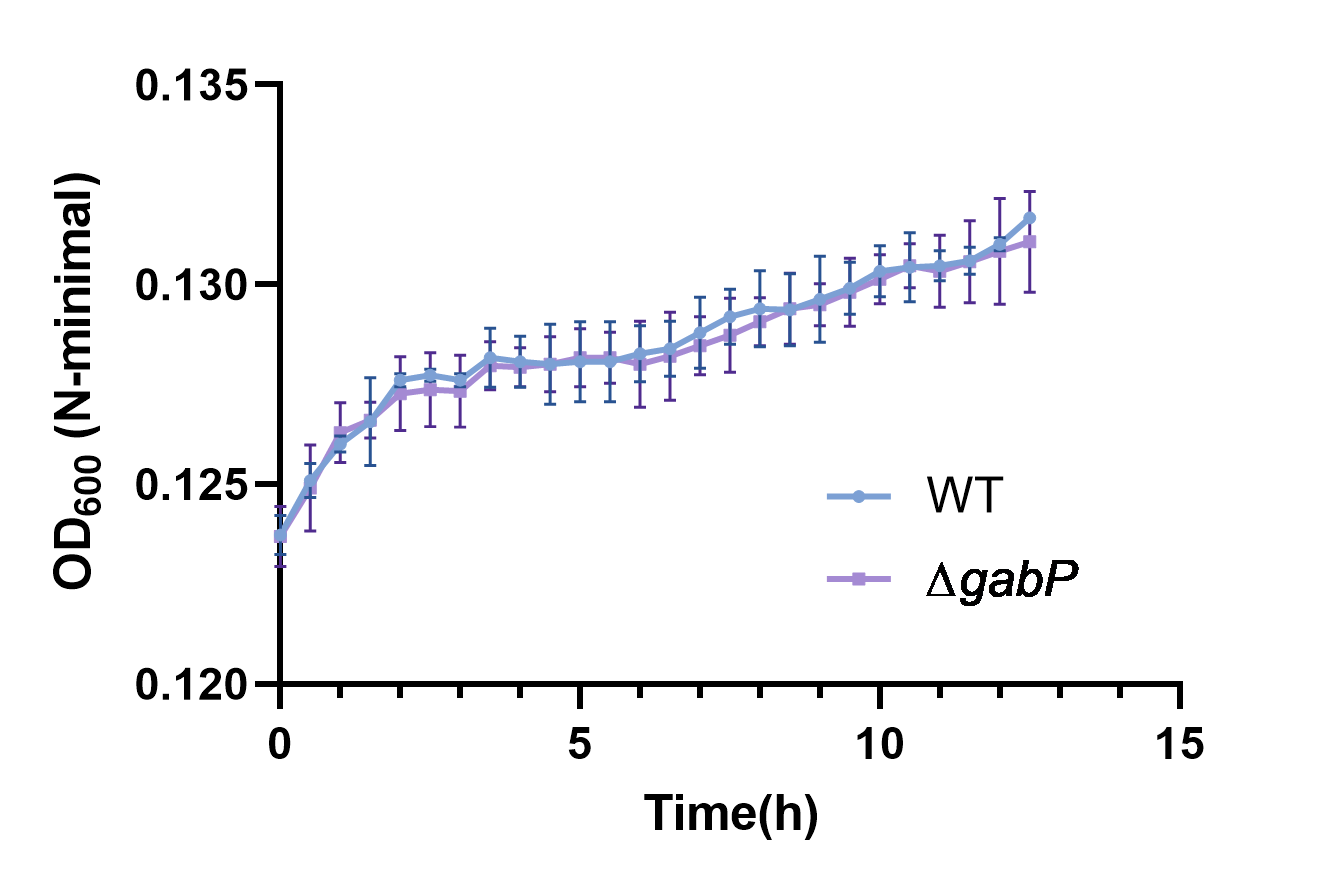

### Figure 1_figure supplement 8

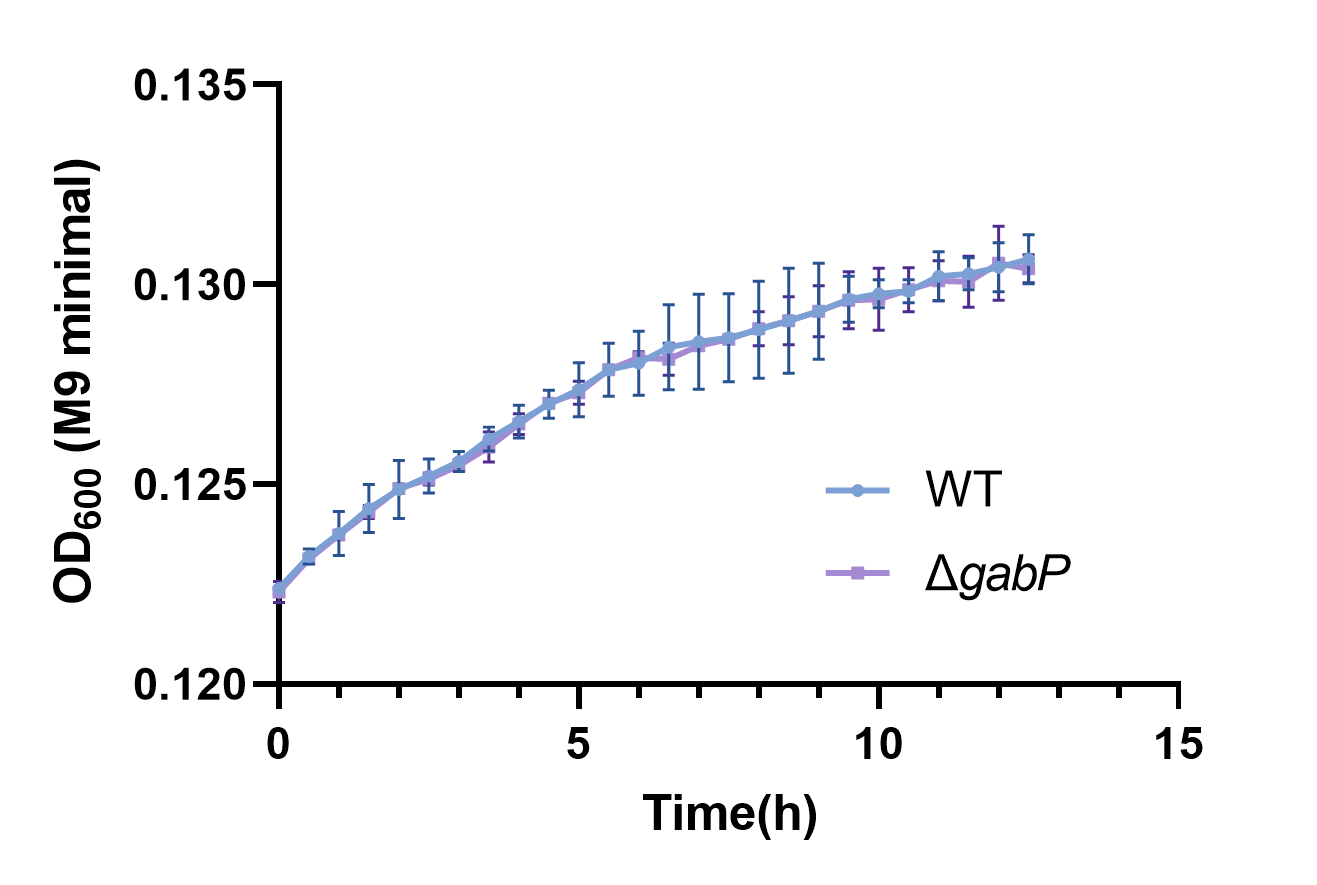

### Figure 2_figure supplement 1

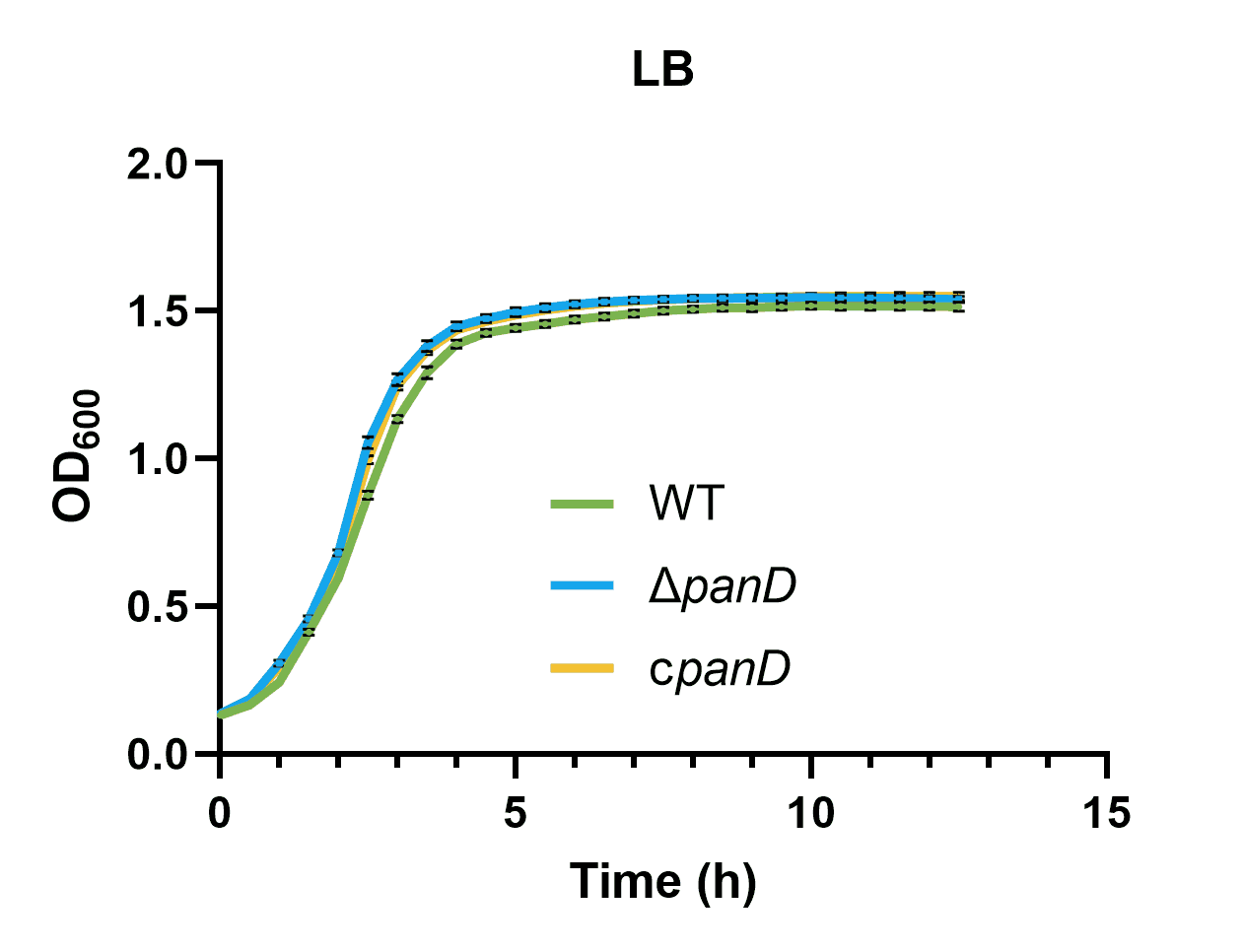

### Figure 2_figure supplement 2

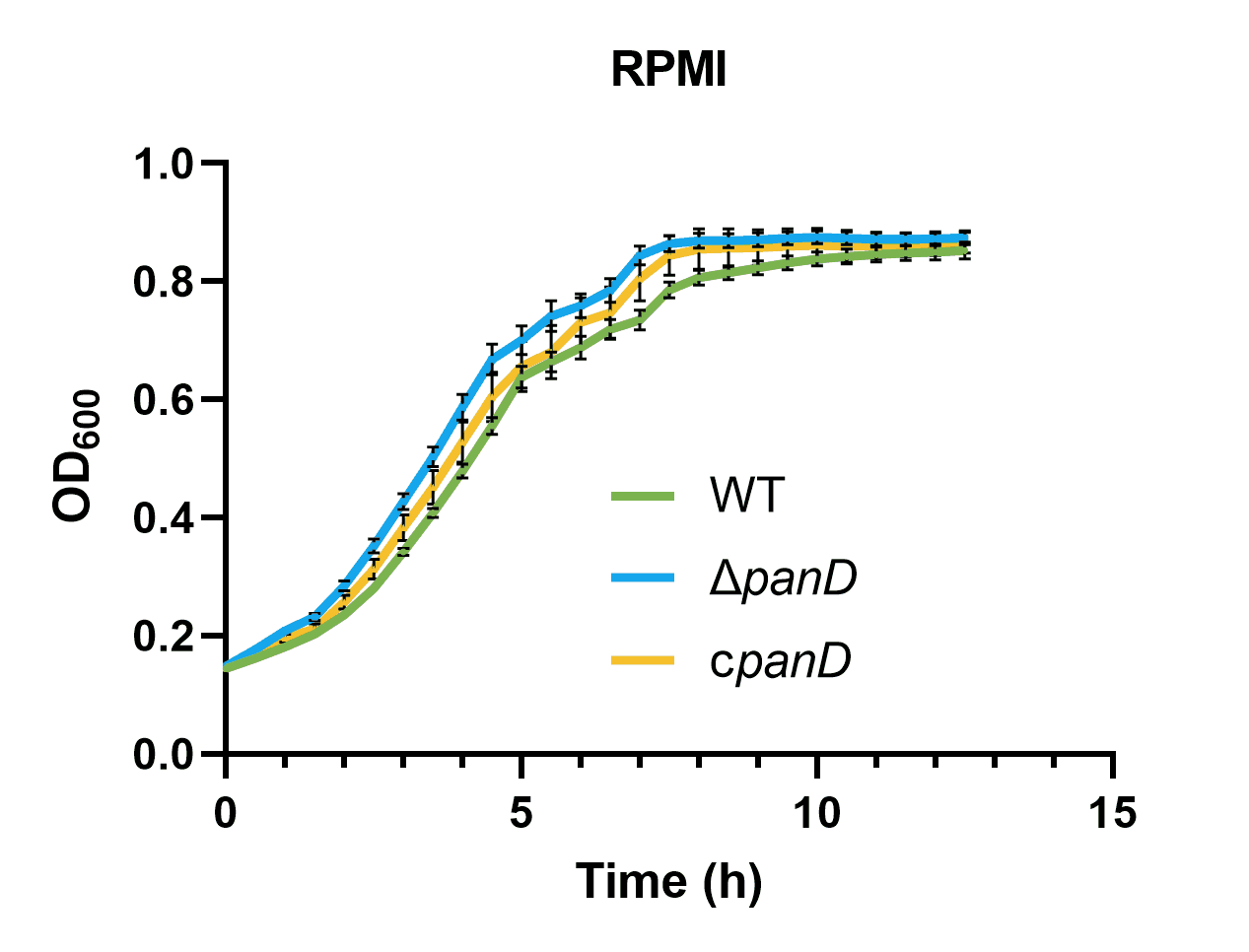

### Figure 2_figure supplement 3

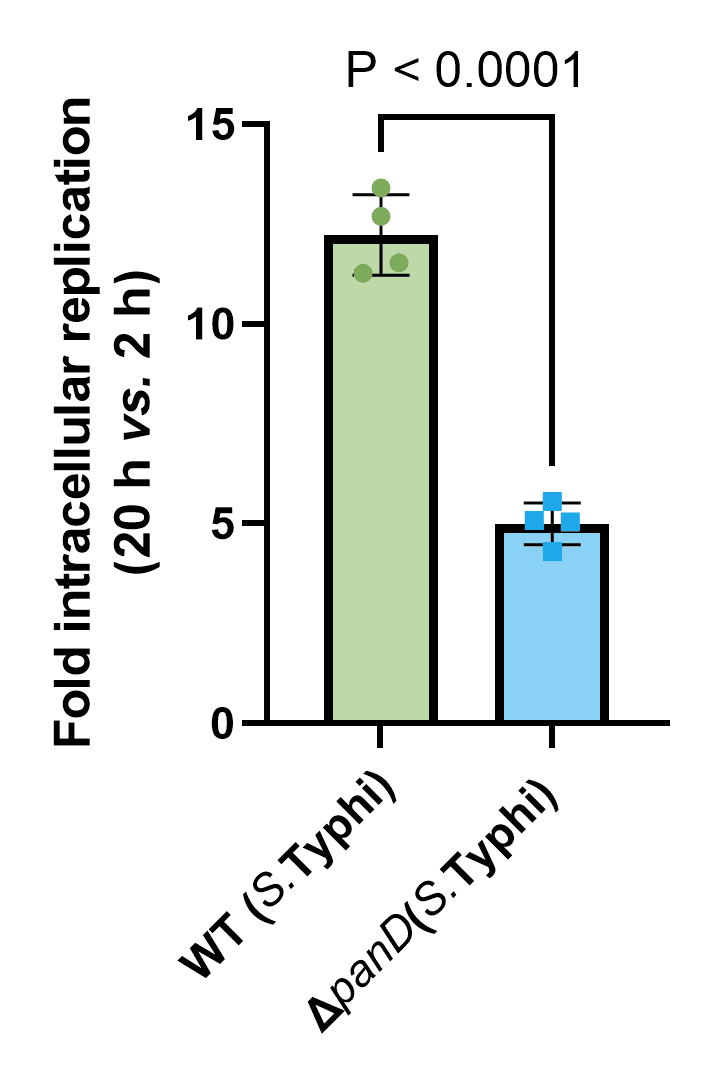

### Figure 4_figure supplement 1

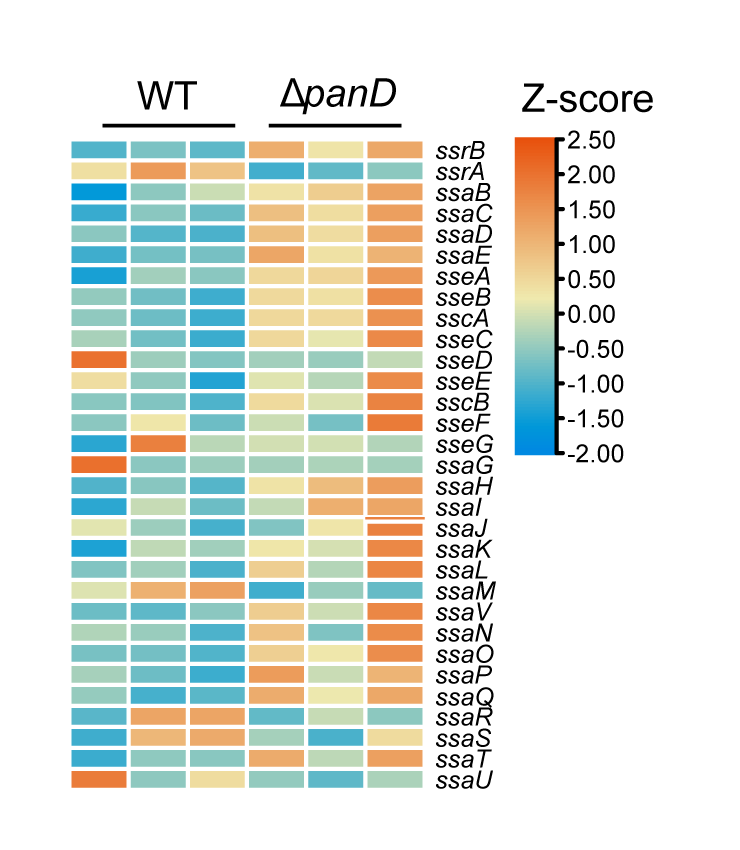

### Figure 4_figure supplement 2

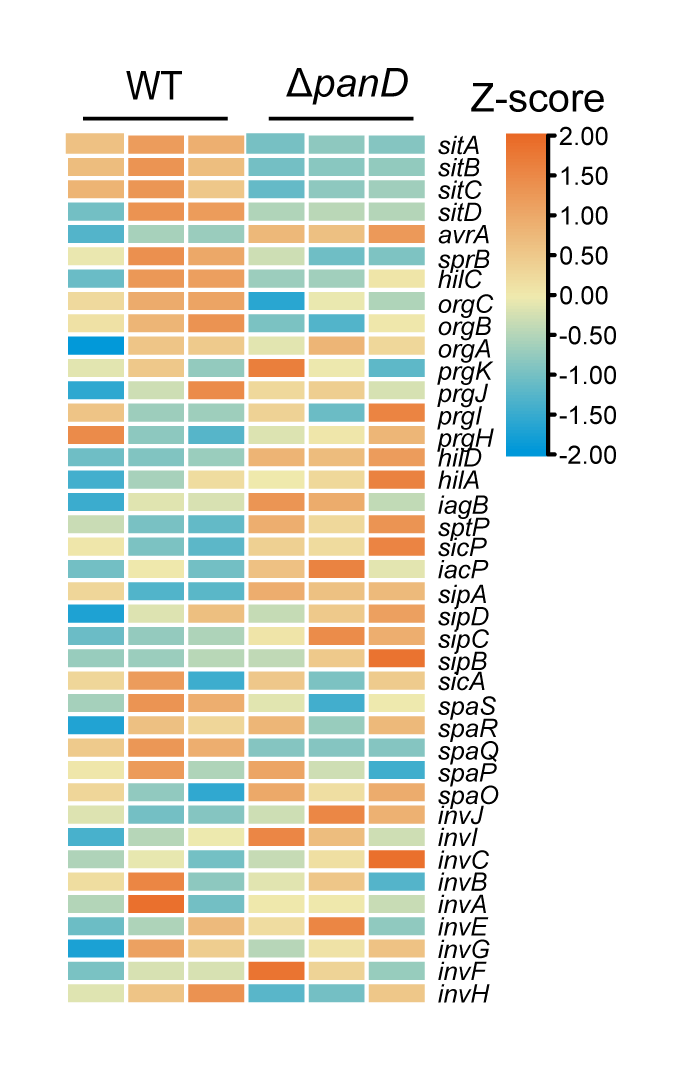

### Figure 4_figure supplement 3

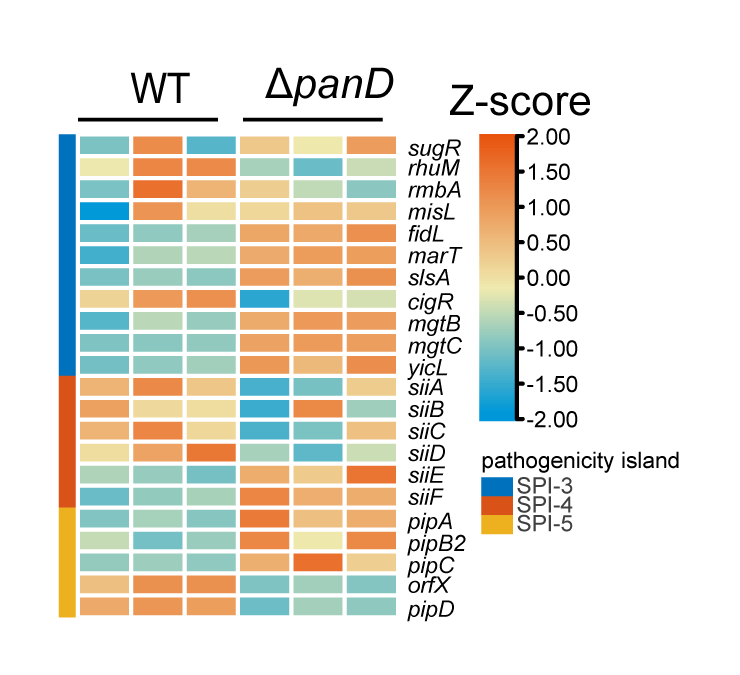
